# Chemically mediated neural and behavioral responses in early benthic juvenile Caribbean spiny lobsters, *Panulirus argus*

**DOI:** 10.64898/2026.01.20.700623

**Authors:** Yuriy V. Bobkov, J. Rudi Strickler, Charles D. Derby

**Affiliations:** Whitney Laboratory for Marine Bioscience, University of Florida, St Augustine, FL, USA; University of Texas Marine Science Institute, Port Aransas, TX, USA; Department of Biological Sciences, University of Wisconsin Milwaukee, Milwaukee, WI, USA; Neuroscience Institute, Georgia State University, Atlanta, GA, USA

**Keywords:** behavior, calcium imaging, chemoreception, Crustacea, lobster, olfaction, olfactory receptor neurons

## Abstract

Spiny lobsters use their chemical senses to acquire resources such as shelter and food, avoid predators, and interact with conspecifics. However, little is known about if and how these responses change over developmental stages. Here, we used early benthic juvenile stage Caribbean spiny lobsters, *Panulirus argus*, in calcium imaging studies to investigate physiological properties of olfactory receptor neurons in the olfactory organ, *i.e*., the antennules, and in behavioral studies to characterize chemically triggered responses. The basic structural organization of the antennules is similar in early benthic juvenile, older juvenile, and adult lobsters. Our calcium imaging studies show that the olfactory receptor neurons of both life stages have generally similar patterns of spontaneous activity, tuning characteristics, sensitivity, and kinetic parameters of responses to chemicals. Our behavioral studies show that early benthic juvenile spiny lobsters have similar behaviors to adults in that they produce currents following stimulation with food-related chemicals, navigate through the chemical plumes to locate the source of food-related chemicals, show alarm responses to conspecific hemolymph, and groom their antennules following stimulation with L-glutamate. Our findings suggest that features of the olfactory organ and its sensory neurons and the behavioral patterns are generally similar across developmental stages, making early benthic juvenile lobsters a favorable model for studying chemosensory transduction, coding mechanisms, and chemical-driven behaviors. The smaller scale of early benthic juvenile lobsters allows the use of compact, miniature benchtop laboratory setups, offering significant flexibility for medium-throughput basic and applied studies.

## Introduction

Spiny lobsters and other decapod crustaceans have well developed chemical senses that they use to detect and identify various chemicals in diverse contexts. Plumes of food related chemicals cause spiny lobsters to alert, search, and find the source [1–3]. Crustaceans stimulated with food related chemicals can also produce currents of different types and functions [4–9]. Besides food related compounds, adult spiny lobsters can detect alarm cues in the hemolymph of conspecifics [10–14]. Certain chemicals can also evoke antennular grooming behavior in spiny lobsters and other decapod crustaceans [15–19].

The chemical senses underlying these behaviors have been well studied in adult spiny lobsters. Crustaceans have several specialized chemical senses that can be categorized as either olfaction or distributed chemoreception, which differ in the location of sensors on the body, organization of sensilla, central projection patterns of the axons of their chemoreceptor neurons, organization of the central neuropil, and function in chemically mediated behaviors [14,20,21]. The spiny lobster’s sensory organs, including the antennules, antennae, mouthparts, and legs, are shown in Fig 1.

**Fig 1.**
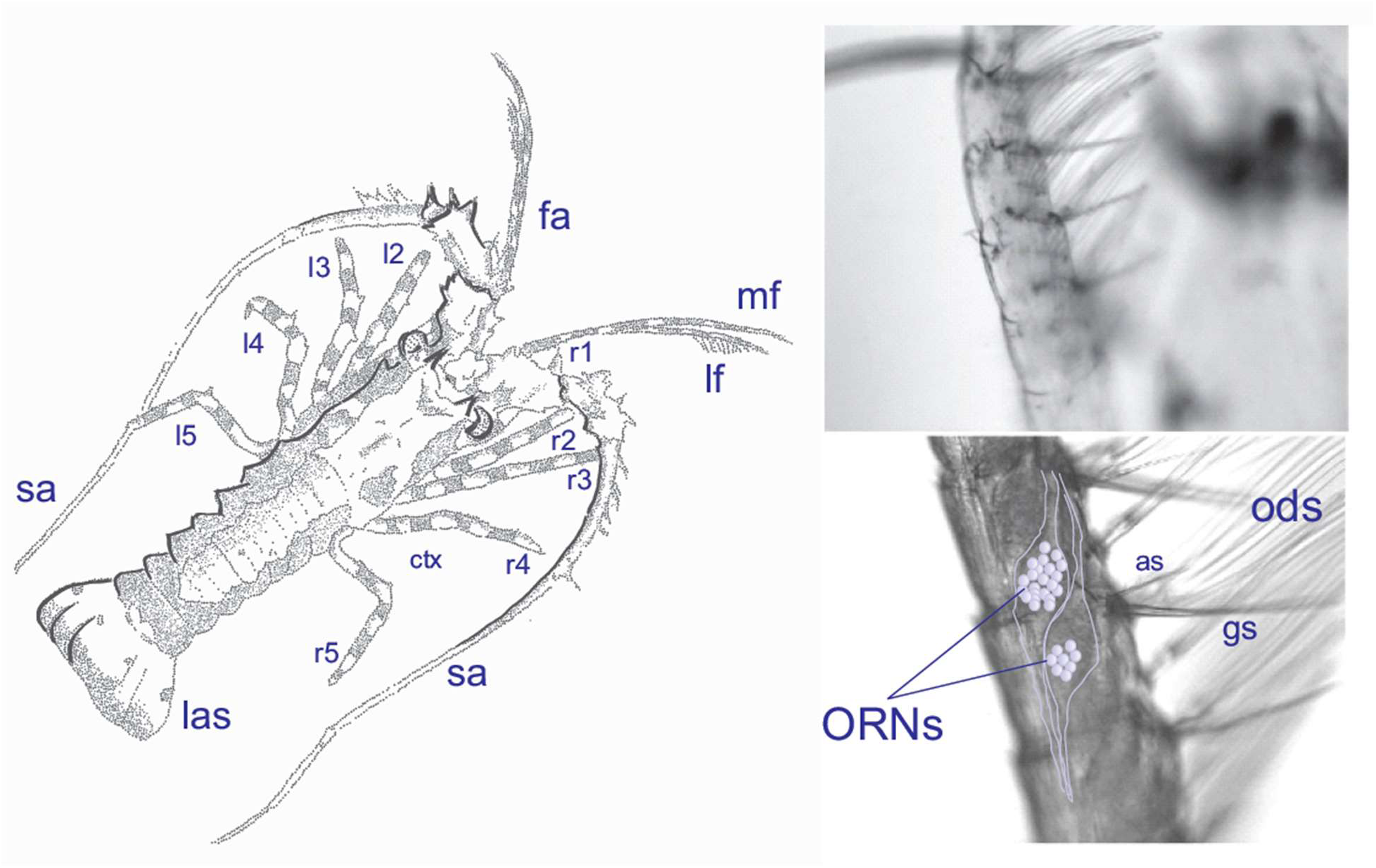
Early benthic juvenile spiny lobsters. <u>Left pane</u>l: lf, lateral flagella of the antennules; mf, medial flagella of the antennules; l1-l5, left walking legs; r1-r5, right walking legs; ctx, cephalothorax; sa, second antenna; abdomen; las, last abdominal segment. Note: pleopods are not visible, as they are on the ventral side of the animal. <u>Right panel</u>: Sensilla on the antennular lateral flagellum. *Top*: Distal region of the lf; *Bottom*: Medial region of the lf. a, aesthetasc sensilla; as, asymmetric sensilla, gs, guard setae, lf, lateral flagellum; ORNs, olfactory receptor neurons; ods, outer dendritic segment of the ORNs.

The olfactory pathway consists of aesthetasc sensilla on the lateral flagella of the antennules, which are innervated by olfactory receptor neurons (ORNs) [20,22]. The olfactory pathway detects both food-related chemicals and pheromones, the latter including hemolymph-based alarm cues, aggregation cues, social cues, and sex pheromones. It mediates orientation toward the source of the cues [23,24]. The second major chemical sense of crustaceans, distributed chemoreception, has a diversity of chemical sensors with properties similar to each other but different from the olfactory pathway in that their sensilla contain both chemosensory neurons and mechanosensory neurons [20]. These bimodal sensilla are located across all body surfaces and appendages. Distributed chemoreception has several subtypes, differing in location of sensilla, central connections, and behaviors that they elicit. For example, distributed chemosensilla on the antennules are like aesthetascs in being sufficient to drive orientation in a chemical plume to find the source of food chemicals. Distributed chemosensory neurons in the asymmetric sensilla, located among other bimodal sensilla on the antennules, uniquely mediate L-glutamate activated antennular grooming behavior [18]. Distributed chemosensory neurons do not mediate responses to pheromones such as alarm cues, aggregation and other social cues, or sex pheromones. Response properties of ORNs and the chemosensory neurons of distributed chemoreception have been examined in adult spiny lobsters, though detailed mechanisms of sensory transduction have been characterized only in ORNs [25–31].

Little is known about the chemical senses and chemosensory behaviors in early stages of the spiny lobster’s life cycle, which is complex [32,33]. It begins with a long planktonic larval phase consisting of phyllosoma larvae with ten stages typically lasting up to 5 to 9 months [34]. The form of phyllosoma larvae is very different from that of early benthic juvenile and adult animals. The phyllosoma larval stages are followed by metamorphosis into the free-swimming, non-feeding puerulus stage, whose body form is much like early benthic juvenile and adult stages though without coloration. Pueruli move into nearshore settlement habitat that includes seagrass, hardbottom, or mangrove habitats [35,36]. Once settled, these early benthic juvenile lobsters begin to feed and are initially solitary. When they reach about 15 mm carapace length, they move from macroalgae to crevice shelters and live there communally for the remainder of their lives. In the Florida Keys, they become adults of legal size at > 76.2 mm carapace length (CL) at ca. 18 months of age since settlement. They have diurnal activity patterns in which they leave shelters at dusk to forage and return to shelters at dawn. Early benthic juvenile spiny lobsters have a body form and olfactory organ that is generally similar form to adults [37,38], though nothing is known about the physiology of their ORNs. Responses of *P. argus* pueruli to chemical cues from red algae associated with settlement and cues from conspecifics have been described [32,39,40]. Responses of early benthic juveniles of other species of spiny lobsters to chemical cues have been described for conspecific cues by *Jasus edwardsii* [41] and habitat cues by *Panulirus cygnus* [42].

Our study examines chemoreception in early benthic juvenile animals. We examined appetitive behaviors to food related cues, antennule grooming behavior to L-glutamate, and alarm responses to body fluids of conspecifics. We also characterized basic properties of the physiology of ORNs. Our results indicate that early benthic juvenile animals are very similar to adults in many of their chemosensory properties.

## Materials and Methods

### Experimental animals

Post-larvae of Caribbean spiny lobsters *Panulirus argus* were collected from floating artificial habitats in the Florida Keys by colleagues at the Florida Fish & Wildlife Conservation Commission in Marathon, Florida Keys [33]. These animals were shipped to our laboratories in Florida and Texas, where they were held in 10 L open glass aquaria (31.8 x 16.5 x 21 cm, TopFin®, Phoenix, AZ, or similar) in groups of 7 to 10 animals. The animals were diet conditioned (food deprived for at least 48 h) before experiments. Aquaria were supplied with flowing natural sea water under local conditions (SW). The early benthic juvenile lobsters used in our study had carapace length (CL) of 6.5 to 9.5 mm (7.8 ± 0.2, n = 20). Some animals were involved in multiple types of experiments. This study was carried out in strict accordance with the recommendations in the Guide for the Care and Use of Laboratory Animals of the National Institutes of Health and according to guidelines at our universities. Animals were chilled on ice for anesthesia or euthanasia, and all efforts were made to minimize suffering.

### Calcium Imaging

#### Preparation

Olfactory receptor neurons (ORNs) were studied using an *in situ* preparation developed earlier [27,29]. Briefly, a single annulus was excised from the lateral flagellum of the antennule, and the cuticle on the side opposite the aesthetasc sensilla was removed to provide better access to the somata of the approximately 200 ORNs associated with each of the approximately 10 aesthetasc sensilla per annulus. Following treatment with trypsin, papain, or collagenase (1 mg/ml, 20 min), the ensheathing tissue covering the clusters of ORNs was gently removed to allow access to somata. All dissections were made in *Panulirus* saline (PS) containing (in mM): 486 NaCl, 5 KCl, 13.6 CaCl_2_, 9.8 MgCl_2_, and 10 HEPES, pH 7.9. The protease cocktail was prepared using divalent cation free PS containing (in mM): 486 NaCl, 5 KCl, 0-10 Glucose, 0.1 EGTA, and 10 HEPES, pH 7.9.

#### Calcium imaging and data analysis

After enzymatic treatment and cleaning, annuli from the antennular lateral flagellum were placed in an Eppendorf tube in PS containing the fluorescent calcium indicator (Fluo-4AM, Invitrogen, Waltham, MA) at 5–10 µM prepared with 0.2–0.06% Pluronic F-127 (Invitrogen, Waltham, MA). The tube was shaken for about 1 h on an orbital shaker (70 rpm). The specimens were transferred into fresh PS, mounted on a glass-bottom 35 mm Petri dishes, and placed on the stage of inverted microscope. Fluorescence imaging was performed on an inverted microscope (Olympus IX-71) equipped with a cooled CCD camera (ORCA R2, Hamamatsu Photonics, Hamamatsu, Shizuoka, Japan) under the control of Imaging Workbench 6 software (INDEC Systems, Santa Clara, CA). FITC filter set (excitation at 510 nm, emission at 530 nm) was used for single-wavelength measurements. Recorded data were stored as image stacks and were analyzed off-line using Imaging Workbench 6 or ImageJ 1.42 (http://rsbweb.nih.gov/ij/index.html). See Fig 2 for an example of fluorescent ORNs and S1 Video and S2 Video for example of spontaneous activity and chemical evoked activity of ORNs. Numerical analyses including event analysis and kinetic parameter estimates were performed using either ClampFit 10 (Molecular Devices, San Jose, CA) or SigmaPlot 10 (Grafiti LLC, Palo Alto, CA). All recordings were performed at room temperature.

**Fig 2.**
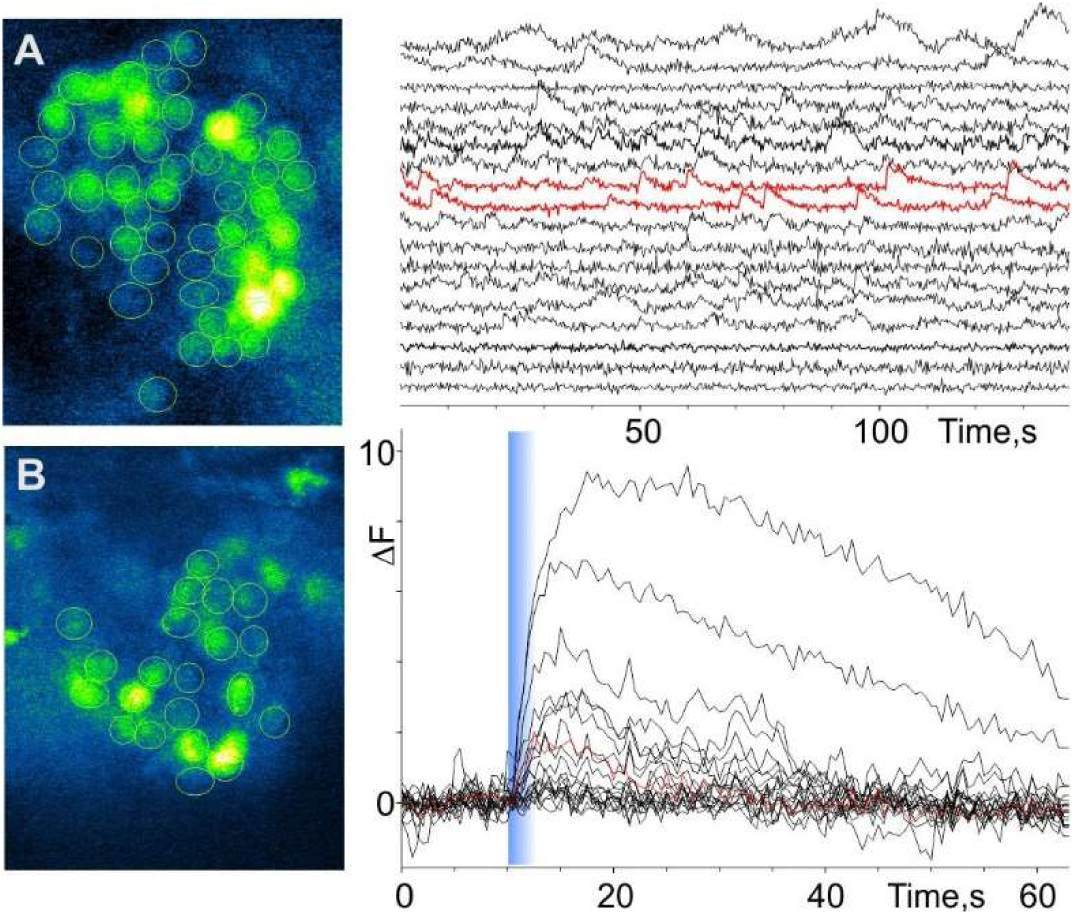
Activity of olfactory receptor neurons (ORNs) from early benthic juvenile spiny lobsters. (A) Spontaneous activity from a cluster of ORNs associated with a single aesthetasc sensillum (*left panel*). The raw calcium signal time series (*right panel*) were background compensated and equally spaced for clarity. Two examples of ORNs characterized by oscillatory activity are highlighted in red. Calcium oscillations are likely associated with bursting ORNs. (B) Chemical evoked activity from a cluster of ORNs associated with a single aesthetasc sensillum different than in A (*left panel*). The *right panel* shows the increase in calcium concentration in ORNs in response to an 800-ms pulse of a chemical stimulus, which was a fish food extract (TetraMarine 0.5 mg/ml). Timing of stimulus presentation is marked by the gray line. Both resting calcium signal level and slow intensity of fluorescence changes caused by bleaching were manually subtracted from each individual trace.

#### Chemical stimuli and their delivery

Unless otherwise noted, the chemical stimulus was an aqueous extract of TetraMarine (TET, Tetra Werke, Melle, Germany), a commercially available marine fish food. Flakes of TET were powdered, dissolved in water (0.1 g/ml), filtered through a 0.2 µm syringe filter, and diluted 1∶200 in PS for experiments. The final maximum concentration was 0.5 mg/ml. The stream containing the chemical stimulus was switched with the flow of PS that otherwise continuously superfused the sensilla (both 250 µl/min) using a multi-channel rapid solution changer (RSC-160, BioLogic, France) under software control (Clampex 10, Molecular Devices, San Jose, CA).

### Behavioral experiments

For observing chemically elicited behaviors, we used two different experimental arenas. Arena type 1 was a relatively large arena that allowed an animal to move over distances and thus allowed us to visualize its chemically activated searching behavior. This arena was either a rectangular glass aquarium with base dimensions of 31.8 cm x 16.5 cm (area of 525 cm^2^) or a Carolina™ culture dish with diameter of 20 cm (area of 314 cm^2^). Both were filled with SW to a depth of 4-5 cm. Animals were introduced into the arena and allowed to adapt for 5-20 min before being exposed to food-related stimuli. In this type of arena, we created a chemical landscape using fragments of shrimp tissue that had been incubated in sea water containing either fluorescein or 6-carboxyfluorescein, 6-CF, (both at 250 μM initial concentration) for at least 2 h. Fluorophores and ambient blue led light (Blue LED Light Panel: 225 psc, λ_em∼465, dimension (cm): 30.5 x 30.5 x 3.2, 1080 Lux, 14W; HQRP, Harrison, NJ) were used to visualize the chemical plume. The animal’s navigation through the chemical plume was recorded on video and analyzed. This set-up is shown in Fig 3.

**Fig 3.**
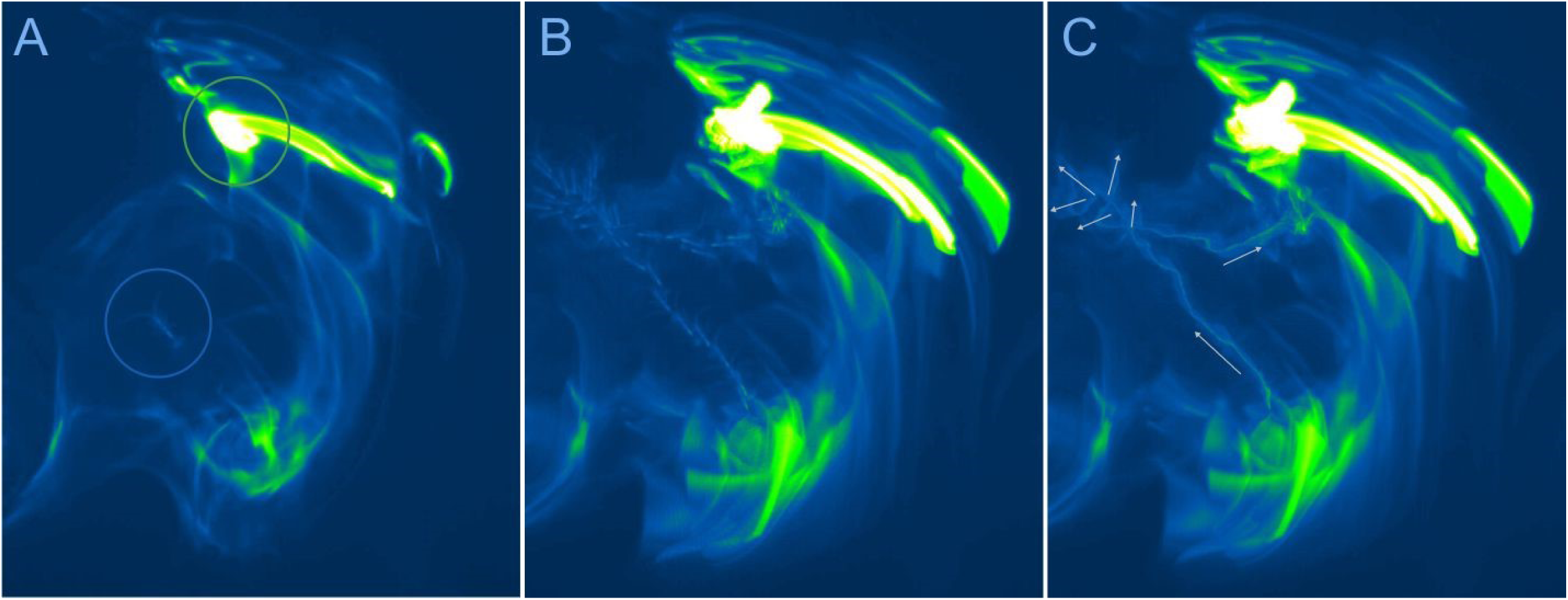
Chemically mediated behaviors of early benthic juvenile spiny lobsters in large (type 1) arena. This arena was a rectangular glass aquarium with base dimensions of 31.8 cm x 16.5 cm (area of 525 cm^2^). This example shows an animal exhibiting search behavior in a food-related (shrimp extract) chemical plume. (A) Single frame of the search trajectory showing the chemical source (*green circle*) and the lobster (marked with blue circle) crossing relatively low intensity portion of the chemical plume. (B and C) The search trajectory is visualized using maximum intensity projection (Fiji/Image J). (B) To better visualize the position of the lobster’s body during search, the number of frames was reduced by a factor of 10. (C) Result of maximum intensity projection of all frames (n = 560). Arrows depict the direction of movement of the animal and vectors of typical exploratory “scent” tracking pattern, cluster of brief error correction moves. See S3 Video for details.

Arena type 2 was smaller and was used for observing and analyzing smaller-scale behaviors triggered by chemical stimuli. This arena was a glass cubic cuvette with dimension (in mm) of either 42.5 x 45 x 45, with optical path length 40; wavelength range: VIS (360-2500 nm), or dimension (in mm) of 52.5 x 55 x 55, with optical path length 50; wavelength range: VIS (360-2500 nm), Helma USA Inc., Plainview, NY. In combination with an aluminized hypotenuse prism, we were able to record simultaneously the top and side views of animals. (The prism was a right-angle prism, N-BK7, 50 mm, λ/8, aluminum hypotenuse, supporting a 425 to 675 nm wavelength range (Newport Corporation, Irvine, CA). This set-up is shown in Fig 4. Animals were video recorded for approximately 2 min in control conditions and then for at least 2 min before, during, and after stimulus presentation. Stimuli were applied as a 5 µl volume delivered using a handheld micropipette to specific body regions such as the antennules or legs. To visualize the plume of the chemical stimulus, we used fluorescein and ambient blue LED light (Blue LED Light Panel: 225 psc, λ_em∼465, Dimension (in mm): 30.5 x 30.5 x 3.2, 1080 Lux, 14W; HQRP, Harrison, NJ).

**Fig 4.**
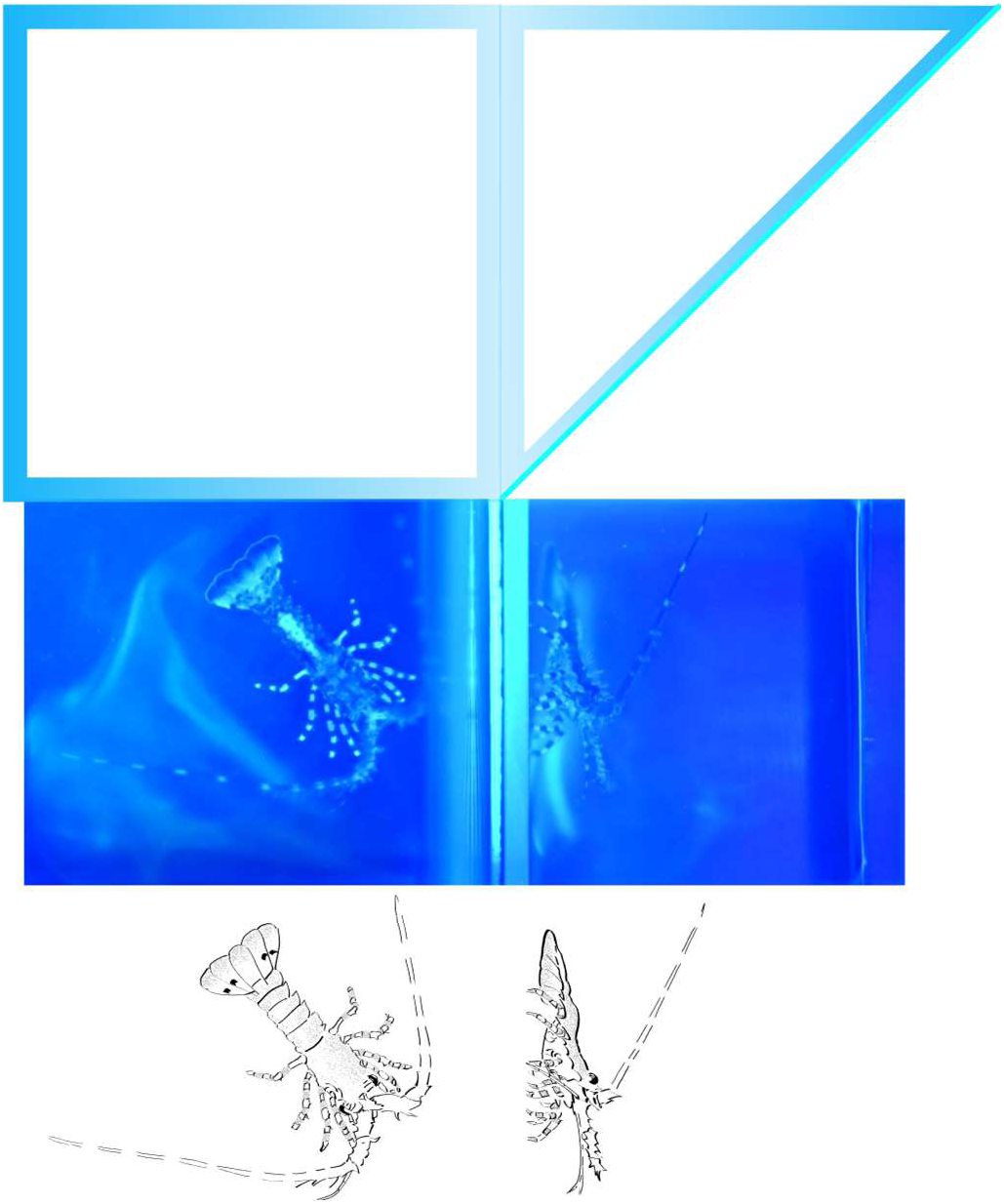
Chemically mediated behaviors of early benthic juvenile spiny lobsters in small (type 2) arena. This arena is a cubic glass cuvette with dimensions (in mm) 42.5 x 45 x 45. A glass prism with an aluminized hypotenuse enabled simultaneous two-side imaging, thus allowing quasi three-dimensional visualization. The animal was exposed to a chemical stimulus (shrimp extract), which was visualized by fluorescein and blue ambient light. This configuration facilitates observing the animal’s sensory appendages including the antennules, antennae, mouthparts, and legs, and how each appendage responds when contacting the chemical plume. <u>Left panel</u>: top view of an animal inside the cuvette. <u>Right panel</u>: side view, prism optical path.

#### Chemical stimuli

Chemical stimuli were mixed in SW that contained fluorescein (250μM initial concentration). The concentration of each stimulus as experienced by the animals was less than that injected since there was some dilution. We estimated a 10-20 times dilution based on intensity of fluorophore assuming linear dependence. We used three chemical stimuli, in addition to SW as the control stimulus. 1) An aqueous extract of shrimp muscle was used as a food-related chemical stimulus. This shrimp extract was prepared by crushing the abdomen of fresh, local penaeid shrimp in SW, incubating it for ca. 2 h, then filtering it through a 0.2 µm filter and diluting it to a final concentration of 5 µl/ml. 2) L-Glutamate at 5 mM was used to determine if animals show chemically activated antennular grooming behavior. 3) A conspecific alarm cue was generated by squeezing the hemolymph and other body fluids from cold anaesthetized live spiny lobsters into SW, then filtering through a 0.2 µm filter for a final concentration of 5 µl/ml.

#### Video recordings

Two cameras were used in behavioral experiments. 1) Nikon Z7 camera, Nikon FX-format, backside illumination CMOS sensor, full-frame 4K UHD/30p video footage, frame size used: 1920 x 1080; at ∼30Hz acquisition rate; lens: AF-S Micro NIKKOR 60mm f/2.8G ED with autofocusing (Nikon Corporation, Tokyo, Japan). 2) Sony HDR-XR500, High Definition, Handycam ‘PAL’ Camcorder, CMOS Sensor with Back-Illumination captures 1920 x 1080 HD 6MP video at 30 Hz frequency (Sony Corporation, Tokyo, Japan). Video recordings were saved in MOV or MTS video formats, converted to AVI using a FFmpeg video converter (free and open-source software, https://ffmpeg.org), and analyzed with ImageJ (free public domain software supported by NIH, USA. http://imagej.org).

## Results and Discussion

We characterized the activity of ORNs in early benthic juvenile spiny lobsters using a preparation developed previously for adult animals in which ORNs were loaded with calcium sensitive fluorescent probe, Fluo-4, and stimulated with odorants delivered to the aesthetasc sensilla containing the outer dendrites of the ORNs. The time course of fluorescent intensity of ORNs of early benthic juvenile spiny lobsters suggests that the spontaneous activity is generally characterized by activity patterns similar to those of adult animals. Some ORNs are tonically active, while other ORNs exhibit pronounced calcium oscillations suggesting bursting activity (Fig 2A and S2 Video) [27,29]. Similar to adult spiny lobsters, the majority of ORNs in early benthic juvenile animals are sensitive to a complex food-related stimulus (Fig 2B and S1 Video, compare with an adult ORN activity from Bobkov et al. [27,29]), while a smaller subpopulation of ORNs was sensitive to monomolecular odorants (data not shown). While ORNs of both early benthic juvenile and adult lobsters are characterized by similar patterns of spontaneous activity and kinetic parameters of the responses, their tuning characteristics and sensitivity to either monomolecular or complex chemical stimuli may differ, though a detailed characterization of these differences and their sensitivities to alarm cues and other pheromones will require further additional future studies. In a first set of behavioral experiments, we evaluated whether early benthic juvenile lobsters exhibit chemically driven search behavior to food related chemicals, a behavior that in adults is mediated by either ORNs or chemosensory neurons on the antennules [23]. For this set of experiments, we put individual diet conditioned early benthic juvenile spiny lobsters in a large (type 1) arena filled with sea water. After a 10-20 min acclimation period, we introduced chemical stimuli into the arena and manually created random artificial plumes visualized with fluorophore. In control experiments, when sea water + fluorophore was used as the stimulus, animals showed no or very little searching for the stimulus source (n = 16, Fig 3). This result is consistent with the conclusion that fluorophore itself as well as both excitation (blue ambient light, λ_em∼465) and fluorophore emission light (λ_em∼518) does not trigger searching behavior in early benthic juvenile lobsters. When a food-related stimulus (shrimp extract) was added to the fluorophore, in a majority cases (14 out of 16 experiments), the animals performed search behavior as soon as they detected the stimulus (Fig 3 and S3 Video). This study sets the stage for future studies that combine hydrodynamics, sensory biology, behavior, and locomotion to help understand mechanisms underlying chemically evoking orientation to chemical cues.

In a second set of behavioral experiments, we tested whether early benthic juvenile lobsters show a diversity of chemosensory behaviors beyond food-evoked searching as has been observed in adult animals. For these experiments, we used the smaller (type 2) arena (Fig 4). We put individual early benthic juvenile lobsters into glass cubic cuvettes with inside dimensions of 40 or 50 mm and imaged simultaneously from the top and lateral views providing possibilities for quasi-3D reconstruction (Fig 4). After 3-5 min acclimation to the cuvette conditions and blue ambient illumination/light provided by blue LED strip light, the animals were video recorded for 2 min in control conditions and then for at least 2 min before, during and after stimulus application. We focused on a limited number of relatively well-described and well-defined behavioral phenomena to avoid ambiguous interpretation. These included a series of food related behavioral patterns, reaction to alarm cues, chemically evoked production of currents, and antennular grooming behavior.

As a food related stimulus, we added 5 µl of shrimp extract in fluorescein into the cuvette using a hand-held pipetter in close proximity to the animal’s head. Fluorescein helped to visualize the plume structure and detection and reaction timings. In all cases (n = 11), the early benthic juveniles were relatively calm in control conditions, exhibited relatively low frequency of antennule flicking and little movement such as walking (Fig 5 left panel, and S4 Video). When food stimulus was presented to the antennules, animals typically increased waving and flicking of antennules and probing with their legs (10 out of 11 trials, Fig 5 right panel and S5 Video). Presentation of the food stimulus to their legs typically resulted in extensive probing with legs, digging, and grabbing at the stimulus.

**Fig 5.**
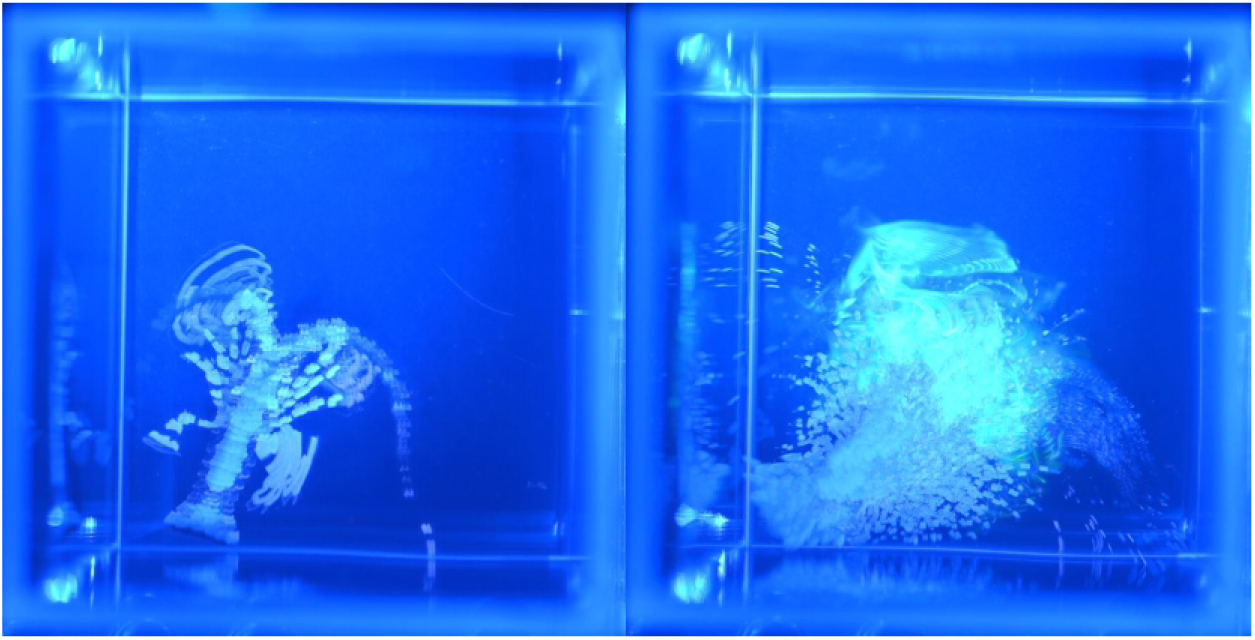
Early juvenile spiny lobster in type 2 arena (small cubic glass cuvette) showing localized searching behavior to a food related chemical stimulus (shrimp extract). <u>Left panel:</u> control (sea water). <u>Right panel</u>: after application of shrimp extract. Activity of juvenile lobster was visualized using maximum intensity projection technique (Fiji/Image J). Note that more articulated body counter suggests minimal movement, compare left and right panels (before and after stimulus application, respectively). The same time interval was used in both cases.

Another behavioral pattern linked to stimulation of chemosensory receptors has been demonstrated in adults: generation of directional water currents. These currents have been previously described for many crustaceans [4–8], including adult spiny lobsters [9], but this has not been described in early benthic juvenile spiny lobsters. These currents can be generated by gills, fan organs (exopodites of mouthparts: crayfish [6,7,43], clawed lobsters [4], snapping shrimp [44], and hermit crabs [45]), abdominal swimmerets (clawed lobsters [4,5]), and paddles on last pair of legs (blue crabs [8]). In crayfish, gill currents are produced by drawing water into the gills from around the animal and expelling it from the frontal opening of the gill chambers using the gill bailers within the gills. It is always directed forward. Fan currents are generated by fan organs associated with the mouthparts (maxillae and maxillipeds) at the anterior end of the animal. These can be directed forward, laterally, upward, and backward [6,7]. In some cases, these currents are described in the context of intraspecific sexual or social interactions that are not related to food. These currents can direct urine released from the nephropores of one animal, the sender, towards another animal, the receiver. They can be used in the context of intraspecific interactions, either reproduction (*e.g*., courtship, mating) or social interactions (*e.g.*, aggression) [4]. Information currents are another current type, being used for both dispersal and reception of chemicals. These currents have been previously described in adult crustaceans. The results of our study show that early benthic stage juvenile spiny lobsters also produce chemically evoked water currents in response to shrimp extract (Fig 6 and S6 Video). In some experiments, when shrimp extract applied directly to the legs, the animals generated currents in a manner similar to the adult animals. While detailed mechanisms triggering these behavioral patterns or the types of currents are not yet described in these early benthic juvenile animals, it is clear that the production of these currents is chemically evoked and conserved across different ontogenetic stages.

**Fig 6.**
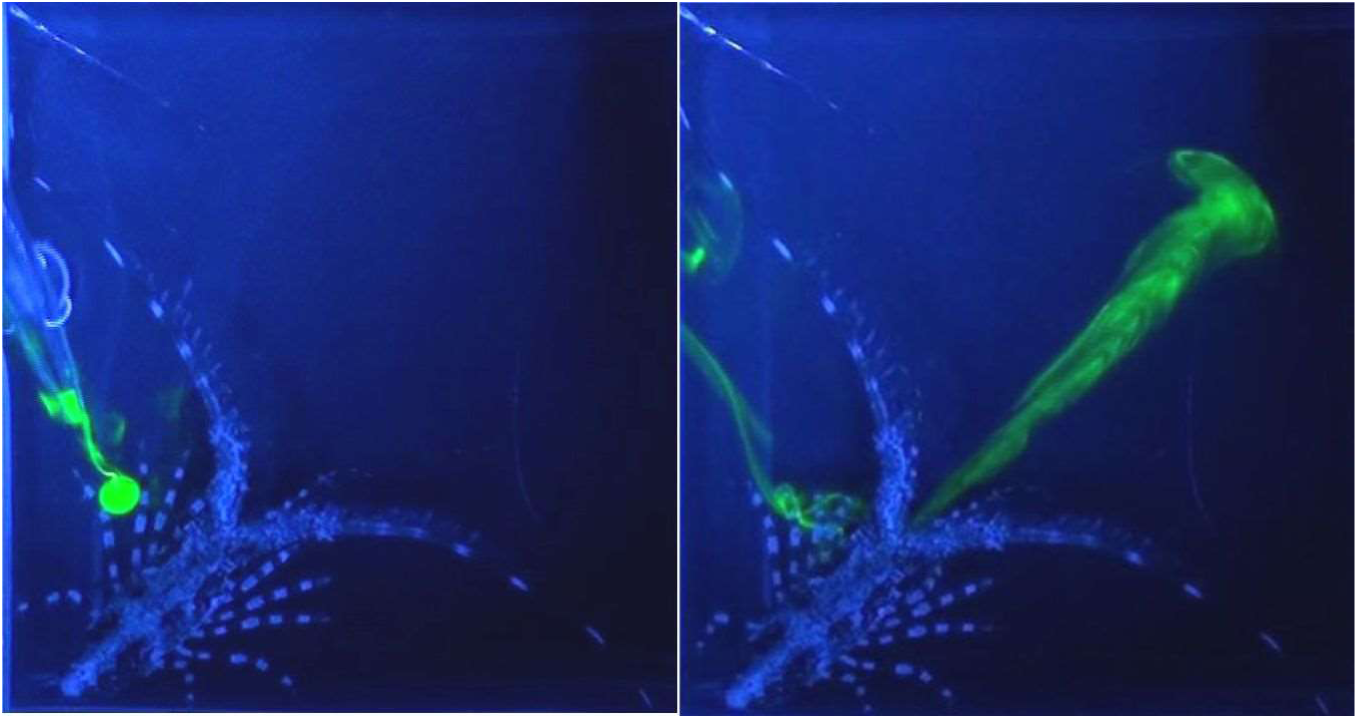
Early benthic juvenile spiny lobster in type 2 arena (small cubic glass cuvette) generating a water current in response to a food-related stimulus. (A) A food related chemical stimulus (5 µl of shrimp extract) was applied to the lobster’s legs using a micropipette. (B) This triggered the animal to generate a current. (B) The animal’s reaction was to generate of current blowing the stimulus away from the anterior end of the animals.

Antennular grooming behavior is one of the most extensively studied and well-understood behavioral patterns, having been described in many decapod crustaceans. It can occur spontaneously (*i.e*., without any apparent sensory input), but it occurs much more often after an animal has been feeding [15–18]. Antennular grooming behavior consists of two major components [18]. In the first, called ‘antennule wiping’, both antennular flagella are repeatedly brought down towards the last pair of mouthpart appendages (third maxillipeds) and are grabbed by their endopodites equipped with pad-like structures consisting of densely packed specialized setae and are repeatedly pulled through these pads. In the second component of antennular grooming behavior, called auto grooming, the two third maxillipeds are rubbed against each other; this usually occurs after a bout of wipes. Antennular grooming behavior can be evoked by relatively high concentration of L-glutamate [17,19,46]. This chemically evoked antennular grooming behavior has been shown only in adult spiny lobsters, and in them it is mediated exclusively by chemosensory neurons in the asymmetric sensilla on the antennules [18]. We therefore examined whether glutamate is capable of evoking grooming behavior in early benthic juvenile lobsters. The response to a 5 µl solution of 5 mM glutamate and fluorescein was initially somewhat similar to reaction to shrimp extract including increased frequency of antennule flicking, antennule waving, and leg probing, but in 8 out of 8 cases, this initial arousal was followed by and/or interrupted by pronounced antennular grooming (Fig 7 and S7 Video). The side view demonstrates this behavioral pattern particularly well. Stimulation with L-glutamate also evoked eye grooming in some cases (S8 Video).

**Fig 7.**
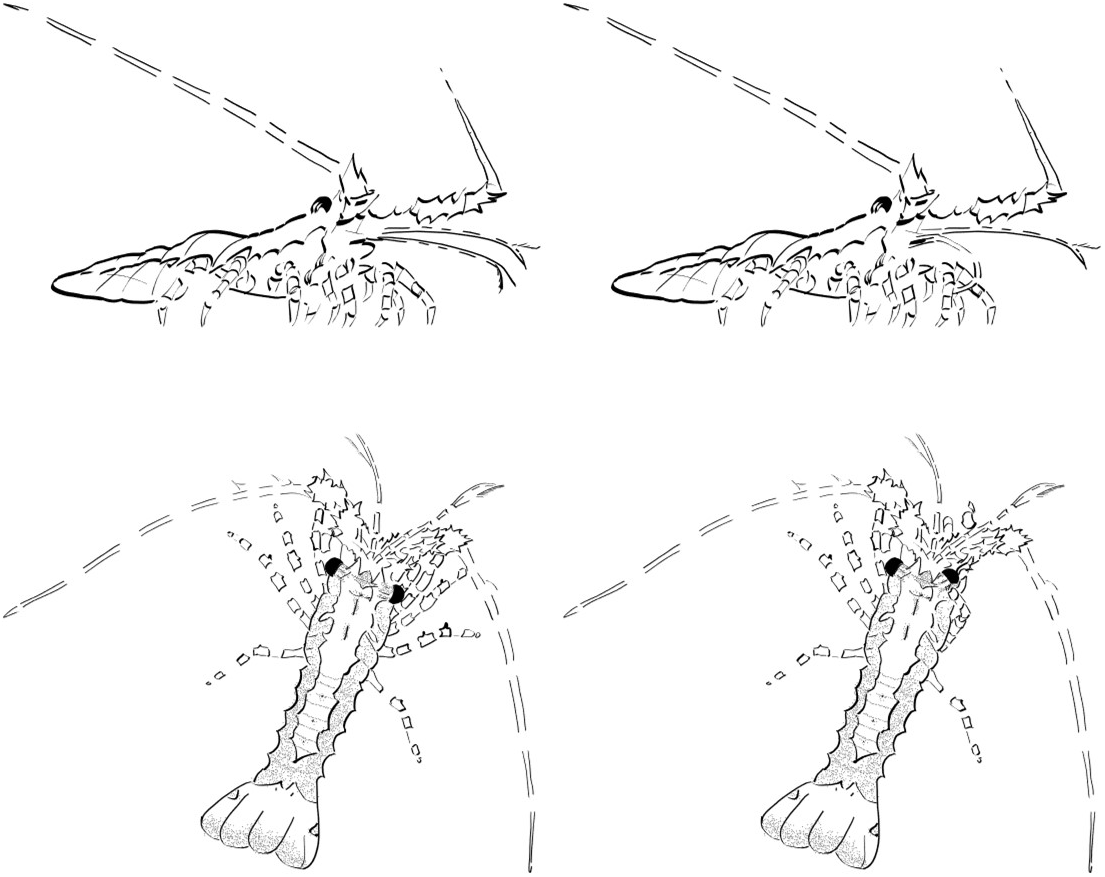
Early benthic juvenile spiny lobster in type 2 arena (small cubic glass cuvette) producing antennular grooming behavior in response to L-glutamate (5 mM, initial concentration). <u>Left</u>: control before glutamate application. <u>Right</u>: after application of L-glutamate. <u>Top</u>: antennular grooming. <u>Bottom</u>: eye grooming.

Another chemically evoked behavior is the response to chemicals released by injured conspecifics, called alarm cues. This has been shown in older juvenile and adult spiny lobsters [10–12]. These alarm cues are present in hemolymph and are detected by ORNs in the antennules of conspecific animals [13]. The molecular nature of the alarm cues in hemolymph has been identified for some adult crustacean species [14] and candidate molecules have been identified in adult spiny lobsters [47]. Behavioral responses to hemolymph-based alarm cues in early benthic stage juvenile spiny lobsters have not been reported. To test whether this stage spiny lobsters exhibit aversive responses to blood-borne conspecific alarm cues, we used the following experimental paradigm: The animals were first recorded in control conditions (Fig 8, S9 Video) and then exposed to food related chemicals (Fig 8, S10 Video). After showing stimulus evoked behavioral excitation, animals were exposed to 5 µl of hemolymph and other body fluids squeezed from conspecifics diluted in SW (Fig 8, S11 Video). At least in 3 out of 4 cases, we observed robust cessation of movement (Fig 8, S11 Video). During this freezing response, the animals lowered their body, stopped walking, and stopped moving their appendages. Aversive responses in general might be characterized by a variety of context dependent behavioral patterns including escape, hiding, aggressive responses and overall hyperactivity. However, the freezing response is one of the most conserved and universal behavioral patterns inherent to a variety of species across different phyla [48,49]. Experimentally, the significant advantage of the freezing response is that it is often a slow event that can be unambiguously detected, especially in space-limited experimental cuvettes.

**Fig 8.**
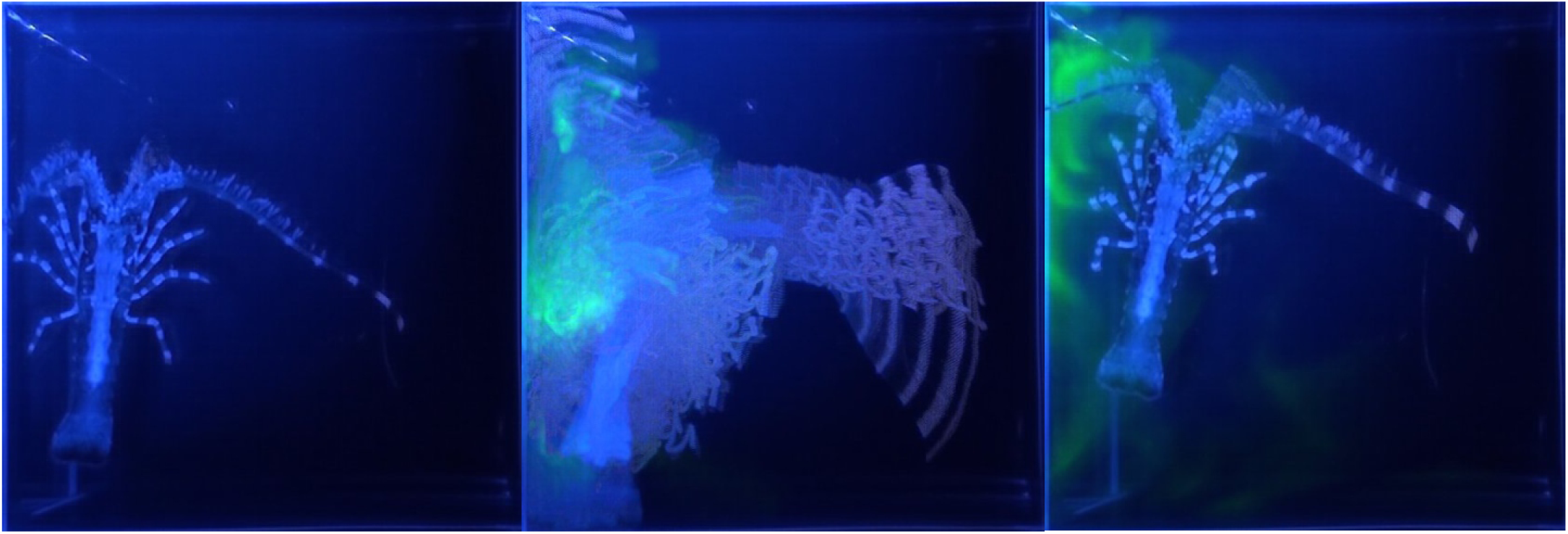
Early juvenile spiny lobster in type 2 arena (small cubic glass cuvette) responding to an alarm cue. <u>Left</u>: negative control (sea water). <u>Middle</u>: food control (shrimp extract). <u>Right</u>: alarm cue (body fluid from a crushed conspecific). Activity of juvenile lobster was visualized using maximum intensity projection technique (Fiji/Image J). Note that more articulated body counter suggests minimal movement. The same time interval was used in all cases.

## Conclusions

Our findings show that the properties of the chemical senses and behavioral patterns of early benthic juvenile spiny lobsters are generally similar to those of adult animals. This makes early benthic juveniles a useful model for studying mechanisms underlying neural activation (*e.g.*, chemosensory transduction and coding) and behavioral responses. The smaller size of early benthic juvenile lobsters confers advantages including the ability to use of a compact, miniature benchtop laboratory setup to rapidly screen candidate compounds from metabolomics and/or bioassay-guided fractionation to identify ecologically relevant chemicals such as alarm cues, sex pheromones, aggregation cues, and food-related cues. In addition, the ability to reliably control conditions within miniature experimental cuvettes and arenas makes it practical to study various environmental factors (including light, visual stimuli, temperature, pH, toxic plankton metabolites, and other favorable or stress conditions) on lobster chemosensory abilities.

---

**S1 Video.** Supporting Fig 2: Calcium imaging of food stimulus evoked activity of ORNs

**S2 Video.** Supporting Fig 2: Calcium imaging of spontaneous activity of ORNs

**S3 Video.** Supporting Fig 3: Searching behavior to food stimulus in large arena

**S4 Video.** Supporting Fig 5: Behavior to control stimulus in small arena

**S5 Video.** Supporting Fig 5: Behavioral responses to food stimulus in small arena

**S6 Video.** Supporting Fig 6: Generation of water current to food stimulus in small arena

**S7 Video.** Supporting Fig 7: Antennular grooming in response to L-glutamate in small arena

**S8 Video.** Supporting Fig 7: Eye grooming in response to L-glutamate in small arena

**S9 Video.** Supporting Fig 8: Response to control stimulus in small arena

**S10 Video.** Supporting Fig 8: Response to shrimp extract presented after control stimulus in small arena

**S11 Video.** Supporting Fig 8: Response to hemolymph presented after shrimp extract in small arena

## Acknowledgments

We thank Dr. Edward Buskey for providing laboratory space at the University of Texas Marine Science Institute, Casey Butler and Thomas Matthews of the Florida Fish & Wildlife Conservation Commission, Marathon, Florida Keys, for providing early benthic juvenile spiny lobsters, and Dr. Joseph Ryan of the Whitney Laboratory for Marine Bioscience, University of Florida, for his valuable discussions and general support.

